# A Zinc-dependent metalloproteinase in the intracellular adaptation of *Brucella abortus* in macrophages

**DOI:** 10.1101/2020.04.17.046490

**Authors:** Leonardo A. Gómez, Francisco I. Alvarez, Raúl Molina, Rodrigo Soto, Carla Daza-Castro, Manuel Flores, Yrvin León, Angel A. Oñate

## Abstract

*Brucella abortus* is a pathogen that survives in macrophages. Several virulence factors participate in this process, including the open reading frame (ORF) BAB1_0270 codifying of a Zinc-dependent metalloproteinase. Here, its contribution in the process of intracellular adaptation was analyzed by infecting RAW264.7 macrophages with the mutant *B. abortus Δ*270 strain. Results showed that this Zinc-dependent metalloproteinase is a cytoplasmic protein that conforms an operon with a transcriptional regulator, which may constitute a type II toxin-antitoxin system. Functionally, this Zinc-dependent metalloproteinase participated neither in the adherence nor the initial intracellular traffic of *B. abortus* in macrophages. Nevertheless, its deletion significantly increased the co-localization of *B. abortus Δ*270 with phagolysosomal cathepsin D, reducing both its co-localization with calnexin, present in endoplasmic reticulum derived vesicles, and its intracellular replication within macrophages. Besides, *B. abortus Δ*270-infected macrophages produced significantly higher levels of TNF-α, IL-6, CD80 and CD86 than *B. abortus* 2308, even when several genes involved in virulence (*vjbR*, *hutC*, *bvrR*, *virB1*) were up-regulated in this mutant. Finally, its deletion significantly reduced the capacity of *B. abortus Δ*270 to adapt, grow and express several virulence factors under acidic conditions. Based on these results, we discuss the role of this Zinc-dependent metalloproteinase in the regulation of the virulence of this pathogen, concluding that it contributes significantly to the intracellular adaptation of *B. abortus* 2308 during the infection of macrophages.

**Author summary:** *Brucella abortus* is the causative agent of the brucellosis, a highly contagious diseases. A Zinc-dependent metalloproteinase contributes significantly in the intracellular survival. Here, we demonstrate that this metalloproteinase has homology with ImmA/IrrE proteases, which are involved in the bacterial resistance to hostile environment. Furthermore, it conforms a gene pair with a transcriptional regulator, being required by *B. abortus* to escape from phagolysosomes, to achieve the endoplasmic reticulum and replicate within macrophages. Its deletion from *B. abortus* stimulated the macrophages, which produced higher levels of pro-inflammatory cytokines and co-stimulatory proteins. This pathogen showed a reduced ability to adapt and grow under acidic conditions, which would negatively affect its escape from phagolysosomes and consequently, stimulating macrophages. Therefore, this work describes how this Zinc-dependent metalloproteinase significantly contributes in the intracellular adaptation of *B. abortus* 2308 in macrophages.

## Introduction

*Brucella abortus* is a facultative intracellular pathogen and the causative agent of brucellosis, a highly contagious zoonosis transmitted from bovines to humans. This is a globally distributed disease, which is acquired through the ingestion of contaminated foods or by the inhalation of aerosols from secretions of diseased animals [1,2]. Infected individuals display irregular fever, headache and arthralgia; its chronicity, associated to the localization of the pathogen in several organs, can produce arthritis, orchitis, endocarditis, hepatitis or encephalomyelitis [1,3,4]. In animals, it shows a tropism towards the erythritol present in placenta and genital organs, producing abortions in females and infertility in males [4,5]. To reach these organs, *B. abortus* survives within neutrophils, macrophages and dendritic cells, using them as “Trojan horses” to disseminate themselves systemically [6–8]. Within these cells, this pathogen develops endosomal *Brucella* containing vesicles (eBCVs), which interact with early and late endosomes and lysosomes, exposing the bacterium to the activity of proteases, reactive oxygen and nitrogen species (ROS and RNS, respectively) and to low pH [9]. Under this environment, *B. abortus* expresses virulence factors allowing it to escape from phagolysosomes and reach the endoplasmic reticulum exit sites (ERES), where it develops replicative vacuoles (rBCVs) to replicate exponentially. Besides, rBCVs can be engulfed by autophagosome-like structures originating autophagic BCVs (aBCVs), which facilitate its exit from infected cells and thus, to complete its intracellular cycle [9–11].

The intracellular survival the *B. abortus* in macrophages requires the expression of several virulence factors such as the lipopolysaccharide (Br-LPS) or cyclic β-1,2-glucans (CβG) [12–14]. Nevertheless, is the type IV secretion system (T4SS) virB a virulence factor fundamental in the development of rBCVs and to complete the intracellular life cycle of this bacteria into macrophages [10–11]. This secretion system is encoded in the *virB* operon, which is induced under the acidic conditions present in the eBCVs and is regulated transcriptionally, among others, by the vacuolar hijacking *Brucella* regulator (VjbR), *Brucella* luxR-like regulator (BlxR), the two component system BvrR/BvrS or the histidine utilization regulator (HutC) [15]. The expression of the T4SS virB system allows *B. abortus* to translocate several effectors to the cytosol of host’s cells, controlling fundamental aspects of the physiology of these cells [9,10,15,16]. One of the best studied effectors is BtpA, a protein with Toll/Interleukin-1 receptor (TIR) domain, which is codified by the BAB1_0279 open reading frame (ORF) present in the genomic island 3 (GI-3) of *B. abortus* 2308 (*B. abortus* 9-941 BruAb1_0274 and *B. melitensis* 16 M BMEI1674) [17,18]. This protein is involved in diverse functions to evade the host’s immune system by the interfering of the intracellular signaling depending on Toll-like receptor 2 (TLR2) and TLR4, inactivation of the Nuclear Factor-kappa B (NF-κB), inhibition of the secretion of pro-inflammatory cytokines or preventing the maturation of dendritic cells [15,18–22]. Theses virulence factors expressed by *B. abortus* into macrophages it allows suppress the signals activating the interferon gamma (IFN-γ)-producing helper T type 1 (Th1) cells and cytotoxic T cells, which allows it to subvert the protective response of host [23–25].

The GI-3 possesses several ORFs participating in the intracellular survival and replication of *B. abortus* [18,20,26,27]. One of them, the BAB1_0270 (*B. abortus* 9-941 BruAb1_0264 and *B. melitensis* 16 M BMEI1683), is of interest because its deletion reduces the intracellular survival and replication of *B. abortus* 2308 in macrophages and epithelial cells as well as bacterial persistence in the spleen of BALB/c mice two weeks after infection [26]. Furthermore, their sequence or part of this ORF (immunodominant epitopes) has been used to develop DNA vaccines capable to induce Th1-type immunogenicity and to confer variable levels of protection in the BALB/c murine model [28–30]. Although the function of ORF BAB1_0270 in the virulence of this pathogen is unknown, *in silico* analyses shows that it codifies a Zinc-dependent metalloproteinase with homology to metalloproteinases of the ImmA/IrrE family present in *Bacillus subtillis* and *Deinococcus* (*D. radiodurans* and *D. deserti*), respectively [31,33]. This ImmA/IrrE metalloproteases breaks repressor proteins (ImmR/Ddro, respectively) impeding the horizontal transfer of mobile genetic elements (e.g. ICE*Bs*1) or the transcription of genes required for DNA repair and survival under conditions such as ionizing and ultraviolet radiation or mutations induced by mitomycin C [31,33]. Hence, if this Zinc-dependent metalloproteinase of *B. abortus* 2308 possesses a similar function to ImmA/IrrE, this function could be to regulate resistance to intracellular microbicide mechanisms of eukaryotic cells, which would explain the attenuation of the virulence of *B. abortus Δ*270 during infection in *in vitro* and *in vivo* models [26].

Therefore, in this work we analyze the contribution of the Zinc-dependent metalloproteinase codified by the BAB1_0270 ORF in the intracellular adaptation of *B. abortus* strain 2308 during the infection process of RAW264.7 macrophages. In accordance with the evidence, this Zinc-dependent metalloproteinase could play important roles in intracellular survival, participating in the regulation of the virulence of this pathogen; consequently, this work attempts to shed new insights on the mechanisms that regulate *B. abortus* 2308 virulence during its establishment in host cells.

## Results

### Physicochemical parameters, antigenicity, modeled refinement and tertiary structure validation

The antigenicity of the Zinc-dependent metalloproteinase according VaxiJen server was 0.3897. This value classified the hypothetical protein as non-antigenic. However, this value is close to the threshold (0.4). Regarding the physicochemical properties, this metalloproteinase has a molecular weight of 21,023.04 Da with an isoelectric point (pI) of 6.01. The half-life of this protein was more than 10 h in *E. coli* (*in vivo*). The aliphatic index and the GRAVY value were 73.96 and −0.423, respectively, indicating the protein stability in a wide range of temperatures and classifies it as hydrophilic. Moreover, the instability index was 37.89, which classifies the hypothetical protein as stable. Prediction of the 3D structure of the hypothetical protein was developed through comparative models with I-TASSER servers. The results showed the best model with the highest C-score of 0.00 which was selected for further refinement. The secondary structure of the hypothetical protein contains 7 helixes, 4 strands and 12 coils. The best model predicted by I-TASSER was refinement and energy minimized. The final model was validated with ProSa-web, which showed a Z-score of −4.52 (Figure 1A) that placed the hypothetical protein in same category of native proteins of similar size. The final 3D model is shown in Figure 1B. The purification and western blot visualization of this Zinc-metalloproteinase showed that has an approximate molecular mass of 22,500 Da (similar to what was predicted by *in silico* analysis) (Figure 1C) and based on experimental and Cello v2.5 predictions it is located in the bacterial cytoplasm (4.018 of reliability).

**Fig 1.**
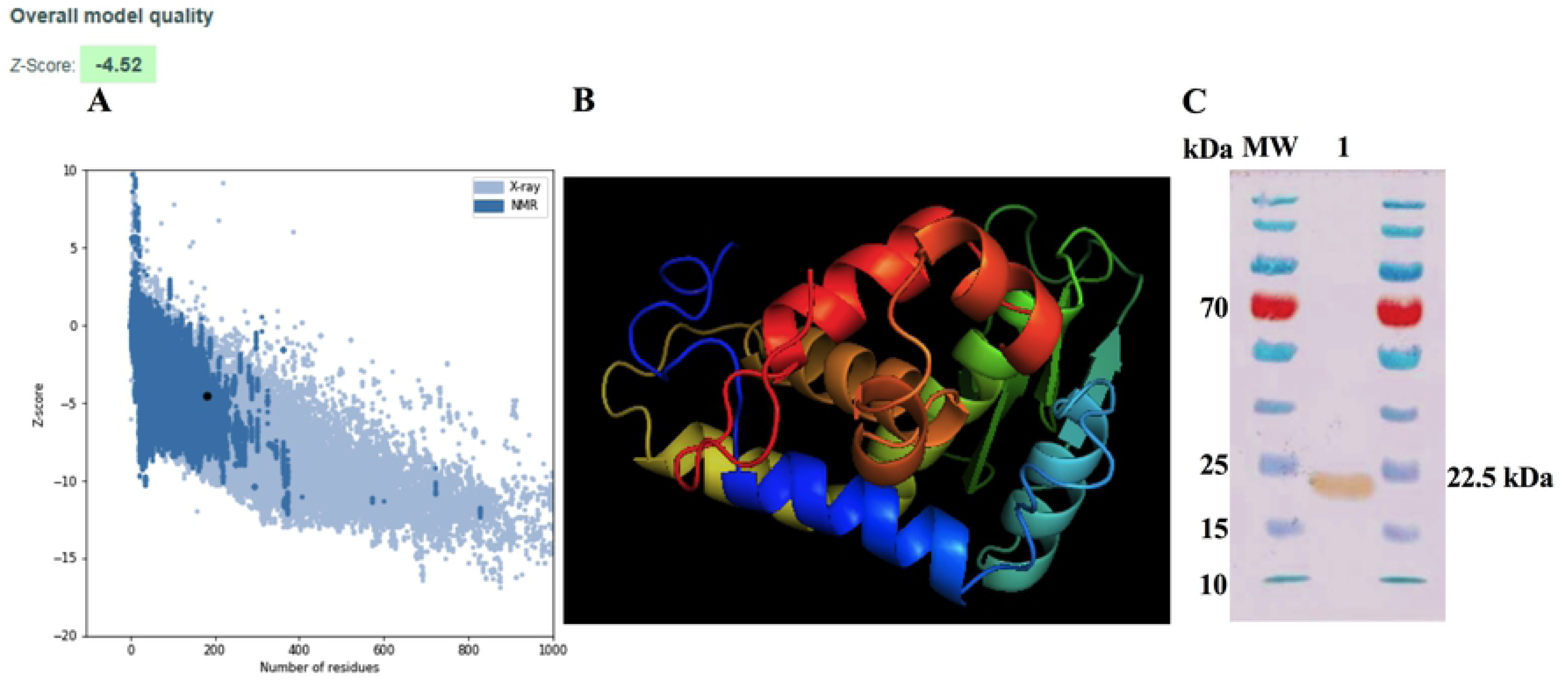
Modelling and purification of Zn-dependent metalloproteinase. **A)** ProSA-web Z-score plot for 3D structure of the hypothetical protein after refinement. The Z-score of the final model is −4.52. The black dot represents the hypothetical protein. **B)** 3D structure according to the I-TASSER server after being refined and visualized by Chimera PyMOL. **C)** Purification of the Zn-dependent metalloproteinase codifying by ORF the BAB1_0270 of *B. abortus* 2308. Zn-dependent metalloproteinase sequences was cloned in the expression vector pColdII (pColdII-Zn-dMP), and recombinant vector was used to transform *Escherichia coli* BL21 (DE3) strain. Western blot was made to visualized protein using anti-6x-HisTag antibodies. MW: molecular weight marker in kDa; 1: recombinant protein purified from *E. coli*.

### A Zn-dependent metalloproteinase that forms part of an operon

The BAB1_0270 ORF possesses a 549 bp nucleotide sequence that encodes this 182 amino acids (aa) Zinc-dependent metalloproteinase. The BLASTp analysis showed that its sequence has homology with the ImmA/IrrE family, metallopeptidases characterized by the presence of conserved COG2856 domains and HEXXH motifs. These metalloproteinases are part of a large family of Zinc-metalloproteinases which form putative operons with proteins having Helix-turn-Helix (HTH) domains of the Xre family. A bioinformatics search demonstrated that this metalloproteinase conforms one unit of transcription (operon) with a transcriptional regulator (WP_002967122.1) localized in the region 270612-271513 of the chromosome I in *B. abortus* 2308 strain (NC_007618.1). These analyses were experimentally confirmed by means of PCR, where the expression of the BAB1_0270 ORF and the transcriptional regulator generated a 906 bp amplicon in the genomic DNA of *B. abortus* 2308 and the same amplicon was obtained from the cDNA (Figure 2A). These results demonstrate that both genes are transcribed together into a single mRNA molecule, where the BAB1_0270 ORF shares four initial nucleotides (ATGA) with the terminal region of gene codifying of a transcriptional regulator, which is constituted by 357 bp and 118 aa, and contains an HTH domain of the Xre family (Figure 2A). Moreover, this operon has a promoter predicted at the −35 and −10 box recognized by RNA polymerase sigma factors rpoD (Figure 2B) and it could be equivalent to the operon codified by the ORFs BruAb1_0264 and BruAb1_0263 of *Brucella abortus* bv. 1 str. 9-941 described by the Toxin-antitoxin Database as a type II toxin-antitoxin system. A hypothetic Type II Toxin-Antitoxin (TA) model for this operon constituted by the Zn-dependent metalloproteinase and the transcriptional regulator is described in the Figure 2C.

**Fig 2.**
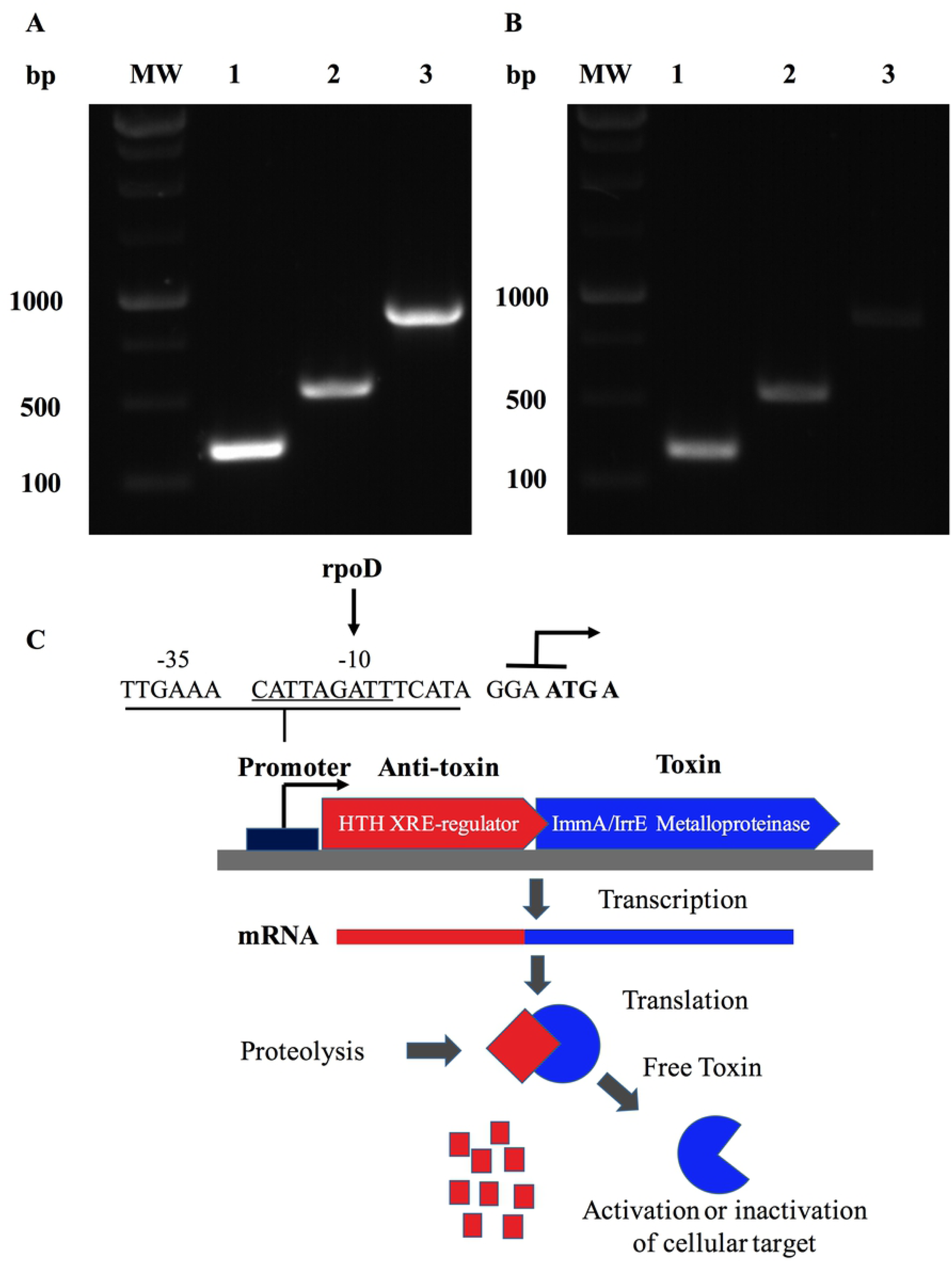
A Zinc-dependent metalloproteinase of *B. abortus* is an operon that forms a putative type II Toxin-Antitoxin. Identification of the transcriptional unit (operon) constituted by the ORF BAB1_0270 and a transcriptional regulator in *B. abortus* 2308 expressed in **A)** the genomic DNA and **B)** the cDNA from total RNA. MW: Molecular weight; lane 1: Transcriptional regulator (357 bp); lane 2: BAB1_0270 (549 bp); and lane 3: operon constituted by ORF BAB1_0270-transcriptional regulator (906 bp). **C)** Hypothetic Type II Toxin-Antitoxin (TA) model for operon constitute by Zn-dependent metalloproteinase and transcriptional regulator. Toxin (Zinc-dependent metalloproteinase) and anti-toxin (transcriptional regulator) are transcribed to mRNA together. Proteases are activated under stress condition, cleaving anti-toxin, which increases the levels of Toxin free, inducing diverse biological functions in bacteria. Predicted promoter at site −35 and −10 binding by RNA polymerase sigma factor rpoD. ATG A: nucleotides share between final part of transcriptional factor and metalloproteinase codified by the BAB1_0270 ORF.

### *B. abortus* adherence and intracellular trafficking into macrophages

The cellular infection by *B. abortus* requires adhesion, invasion and the developed of a replicative niche associated to ER derived vesicles. The role of BAB1_0270 ORF on this process was evaluated infecting RAW264.7 macrophages with *B. abortus Δ*270. This mutant strain showed levels of adherence to macrophages comparable to the wt strain (*P* > 0.05) (Figure 3A and B). Nevertheless, its capacity to invade macrophages, measured by its co-localization with the early endosome marker EEA1, was variable because at 5 min pi the mutant strain showed not significant differences compared to wt strain (*P* > 0.05). However, its percentages of co-localization were significantly lower at 10 min (*P* < 0.001), while at 15 min pi its co-localization was significantly higher than the wt strain (*P* < 0.05) (Figure 3C and D). Later, at 1 h the mutant and wt strains shows similar levels of co-localization with cathepsin D, a phagolysosomal protease (*P* > 0.05) (Figure 5E and F); however, at 12 h pi differences between both strains were observed, where the mutant strain showed significantly higher levels of co-localization with cathepsin D than the wt strain (*P* < 0.01) (Figure 5E and F). Additionally, the co-localization of both strains with calnexin, a chaperone protein present in the ER derived vesicles, was comparable at 1 h pi (*P* > 0.05), but at 12 h pi, the levels of co-localization of the mutant strain were significantly lower than wt (*P* < 0.01) (Figure 5G and H). These results suggest that the deletion of this Zn-dependent metalloproteinase reduces in *B. abortus Δ*270 its capacity to escape from phagolysosomal compartments, would affect significantly its capacity to achieve the ER derived vesicles of macrophages.

**Fig 3.**
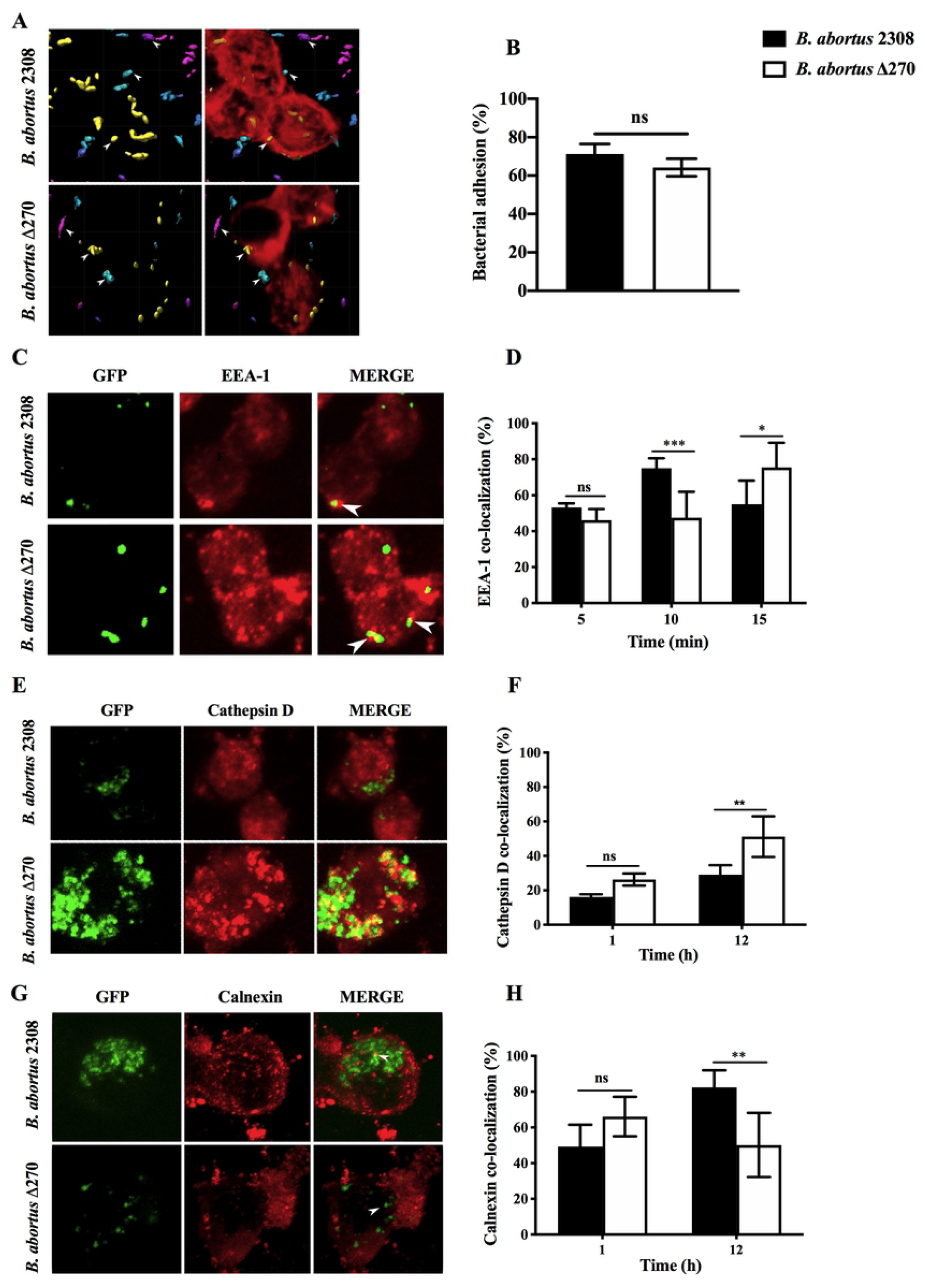
Adhesion and intracellular trafficking of *B. abortus* strains in RAW267.4 macrophages. **A)** Macrophages were treated with cytochalasin D (0.5 mg/ml) and infected with *B. abortus* 2308 and *B. abortus Δ*270 strain for 30 min. Image shows the proximity of *B. abortus* strains with the RAW264.7 macrophages, where cyan represent bacteria with a proximity less to 5 ìm of macrophages; purple represent bacteria with a proximity higher 5 ìm of distance to macrophages, and yellow represent bacteria adhered to macrophages. **B)** Adherence is expressed by the percentages of adhered bacteria (yellow) to RAW264.7 macrophages. Images **C, E and G)** are representatives of confocal images showing the co-localization of *B. abortus*-GFP strains with EEA1 at 5, 10 and 15 min, cathepsin D at 1 and 12 h and calnexin at 1 and 12 h. **D, F and H)** shows the percentages of the co-localization of *B. abortus*-GFP strains with EEA1 at 5, 10- and 15-min pi; cathepsin D at 1 h and 12 h pi, and calnexin at 1 and 12 h pi. Results are expressed as the mean ±standard deviation. Values of *P* < 0.05 were considered as statistically significant, where ns: non-significant differences and *, ** and *** denotes values of *P* < 0.05, *P* < 0.01 and *P* < 0.001, respectively. All assays were made in triplicates.

### *B. abortus* intracellular survival and its gene expression during infection of macrophages

This pathogen replicates in the macrophages by the expression of several virulence factors, including the BAB1_0270 ORF. The role of this ORF in the intracellular survival and the expression of several genes involved in the virulence was evaluated infecting RAW264.7 macrophages. At 6 h pi, the intracellular survival of this mutant strain showed CFU/ml counts significantly lower compared to the wt strain (*P* < 0.001). Besides, the *B. abortus Δ*270 counts were maintained relatively low and constant from 6 h to 24 h, a time where were registered higher differences between the mutant and wt strains (*P* < 0.0001) (Figure 4A). At 24 h pi wt strain was characterized by a robust replication, being higher than the mutant and complemented strains. Additionally, at 24 h pi the gene expression of this mutant strain showed that *vjbR* (*P* < 0.01), *hutC* (*P* < 0.01) and *bvrR* (*P* < 0.05), genes involved in the response to stress (e.g. acidic stress), quorum sensing (QS) or expression of virulence factors (e.g. T4SS virB) were significantly upregulated (Figure 4B). Furthermore, the *virB1* genes showed an up-regulation in this mutant strain, but not differences were observed for *virB2* and *virB5* (Figure 4B). Among the genes codifying of effector proteins translocated through of the T4SS virB (*vceA*, *vceC* and *btpA*), the *vceA* gene showed a down-regulation in *B. abortus Δ*270 compared to the wt strain (*P* > 0.05). Finally, in the mutant strain was downregulated the GI-3 ORF BAB1_0273, codifying of a hypothetical transcriptional regulator, and the ORF BAB1_0627, encoding of a hypothetical protein of unknown functions. These results would demonstrate that deletion of this Zn-dependent metalloproteinase produces, direct or indirectly, changes in the expression of diverse genes involved in the virulence and resistance of this bacteria during infection in macrophages.

**Fig 4.**
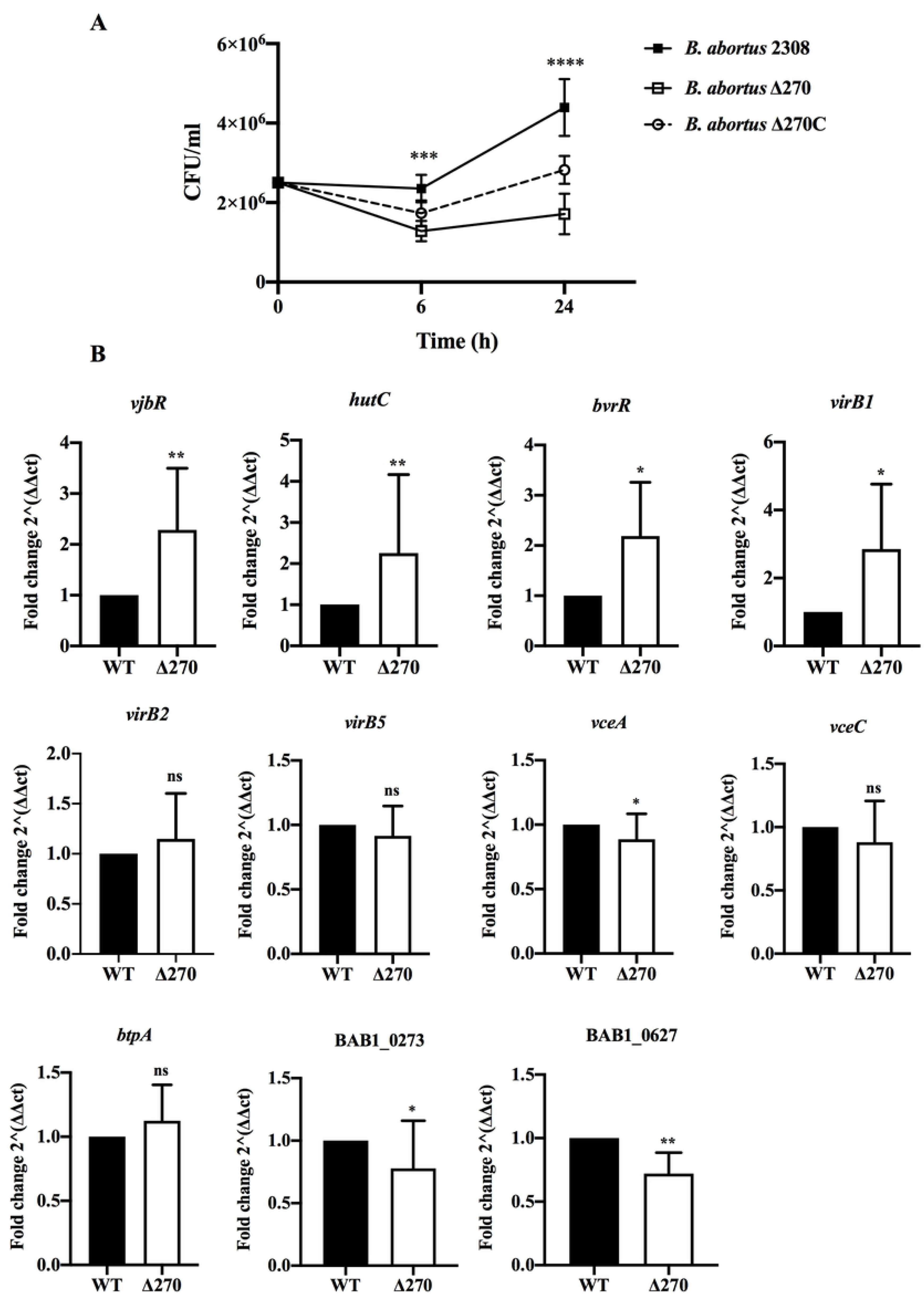
Intracellular survival and gene expression of *B. abortus* strains in macrophages. **A)** Intracellular bacteria were obtained from macrophages infected at 1:10 MOI with *B. abortus* 2308, *B. abortus Δ*270, or *B. abortus Δ*270C at 6 h and 24 h. Intracellular survival was registered as mean ± standard deviation of bacterial counts as CFU/mL. **B)** The relative expression of *vjbR*, *hutC*, *bvrR*, *virB1*, *virb2*, *virB5*, *vceA, vceC, btpA* and the BAB1_0273 (GI-3) and BAB1_0627 ORFs induced in *B. abortus* 2308 (WT) and *B. abortus Δ*270 during infection of macrophages was calculated by the 2-ΔΔ*CT* method using qPCR assays at 24 h pi. The housekeeping *gyrA* and *16s* genes were used as reference genes. Results were expressed as the mean ± standard deviation. Values of *P* < 0.05 were considered as statistically significant, where ns denotes non-significant differences, *, *** and **** denotes values of *P* < 0.05, *P* < 0.001 and *P* < 0.0001, respectively. All assays were made in triplicates.

### Macrophages activated by *B. abortus* strains

Macrophages destroys bacteria in the phagolysosome compartments and activates the innate and adaptive immunity secreting pro-inflammatory cytokines (TNF-α and IL-6) and co-stimulatory proteins (CD80 and CD86), which were quantified at 6 and 24 h by ELISA and flow cytometry, respectively. Macrophages infected with the *B. abortus Δ*270 strain at 6 h pi produced levels of TNF-α significantly higher than the macrophages infected with wt (*P* < 0.001), inactivated strain (*P* < 0.05) and unstimulated macrophages (*P* < 0.0001). At 24 h pi, the macrophages infected with the mutant strain secreted even higher levels of TNF-α compared with different groups (*P* < 0.001) (Figure 5A). Similarly, macrophages infected with the mutant strain at 6 h pi secreted not significant levels of IL-6 compared to wt or inactivated strains (*P* > 0.05), showing statistically significant differences only with non-infected macrophages (*P* < 0.05). However, at 24 h pi, the macrophages infected with *B. abortus Δ*270 produced levels of IL-6 highly significant compared with the production of this cytokine by macrophages infected with wt, inactivated or non-stimulated macrophages (*P* < 0.0001) (Figure 5B). On the other hand, macrophages infected with *B. abortus Δ*270 at 6 h showed not difference in the expression of the CD80 and CD86 proteins compared with macrophages infected with wt, inactivated or non-stimulated (Figure 5C, D and E). However, the expression of these co-stimulatory proteins in macrophages infected with *B. abortus Δ*270 at 24 h were significantly higher compared to wt, inactivated or unstimulated-macrophages (Figure 5C, D and E). These results suggest that the deletion of this Zn-dependent metalloproteinase attenuates this mutant strain, activating the macrophage’s defensive mechanisms required in the clearance of this pathogens.

**Fig 5.**
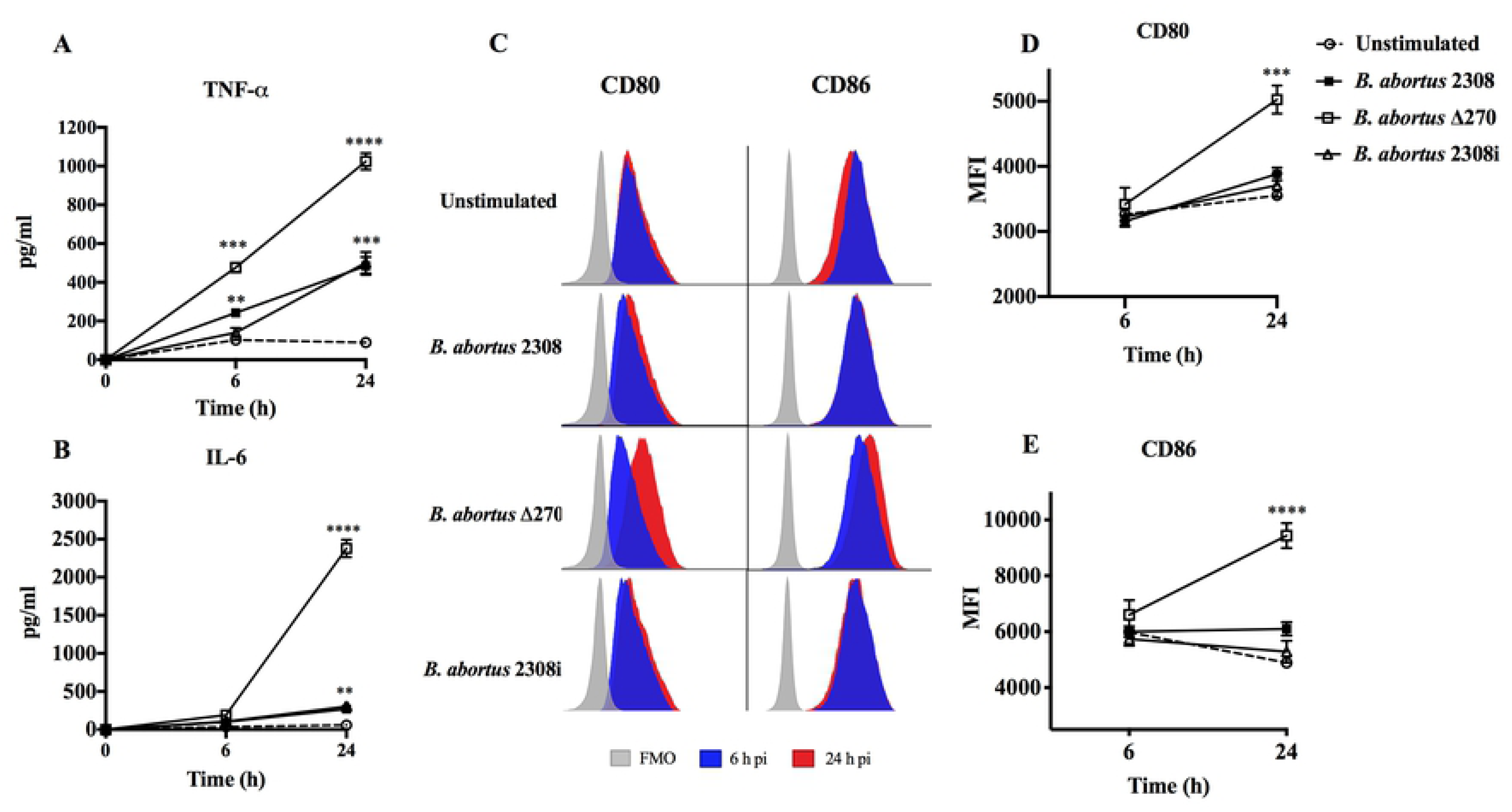
Macrophages activated by *B. abortus* strains. Expression of pro-inflammatory cytokines and co-estimulatory proteins from *B. abortus* strains-infected macrophages at MOI 1:10. **A)** production of TNF-α (pg/ml) and B) production of IL-6 (pg/ml) from infected-macrophages with *Brucella* strains were measured by Sandwich ELISA at 6 h and 24 h pi using standard curves using TNF-α and IL-6 recombinant proteins. **C)** Expression of co-stimulatory proteins CD80 and CD86 from infected-macrophages measured by Flow cytometry at 6 h (blue) and 24 h (red) pi. **D and D)** show the mean ± standard deviation of the fluorescent intensity (MFI) of CD80 and CD86. Data represents unstimulated (PBS treated) macrophages, *B. abortus* 2308-infected macrophages, *B. abortus Ä*270-infected macrophages and *B. abortus* 2308i (inactivated strain) stimulated macrophages. Results were expressed as mean ± standard deviation. Values of *P* < 0.05 were considered as statistically significant, where **, *** and **** denotes values of *P* < 0.01, *P* < 0.001, *P* < 0.0001, respectively. All assays were made in triplicates.

### Bacterial growth and gene expression in acidic conditions

*B. abortus* survives to the acidification produced in the phagosome-lysosome compartments. Here, the contribution of BAB1_0270 ORF in the growing and the gene expression of *B. abortus Δ*270 under acidic conditions was measured. Growth curves of *B. abortus Δ*270 showed a significant lower capacity to adapt and grow in a medium with pH 5.5 compared to wt or complemented strains (*P* < 0.0001) (Figure 6A). Furthermore, this mutant strain was maintained in the lag phase from 6 h until 72 h of culture, entering to logarithmic phase at 72 h (exponential growth). WT and complemented strains showed similar growth curves in these acidic pH conditions, with a lag phase that extends to the 48 h time point, and an exponential phase between the 72 h and 96 h time points. Interestingly, these strains developed different curves of growth in this acidic medium, linear correlation analysis showed that the three strains increased the pH in the medium, showing not significant differences in the pH during the growing of the *Brucella* strains (*P* > 0.05) (Figure 6B). Finally, the relative gene expression of *B. abortus Δ*270 cultured in this medium showed a lower expression of the genes *vjbR*, *hutC*, *bvrR*, *virB1*, *virB2*, *virB5*, *vceA*, *vceC* and *btpA* compared to wt strain (*P* < 0.05) (Figure 6C). These results suggest that deletion of this Zn-dependent metalloproteinase reduces the capacity of *B. abortus* to adapt and grow under acidic conditions and produces, directly or indirectly, changes in the expression of diverse genes involved in its virulence and resistance to acidic conditions.

**Fig 6.**
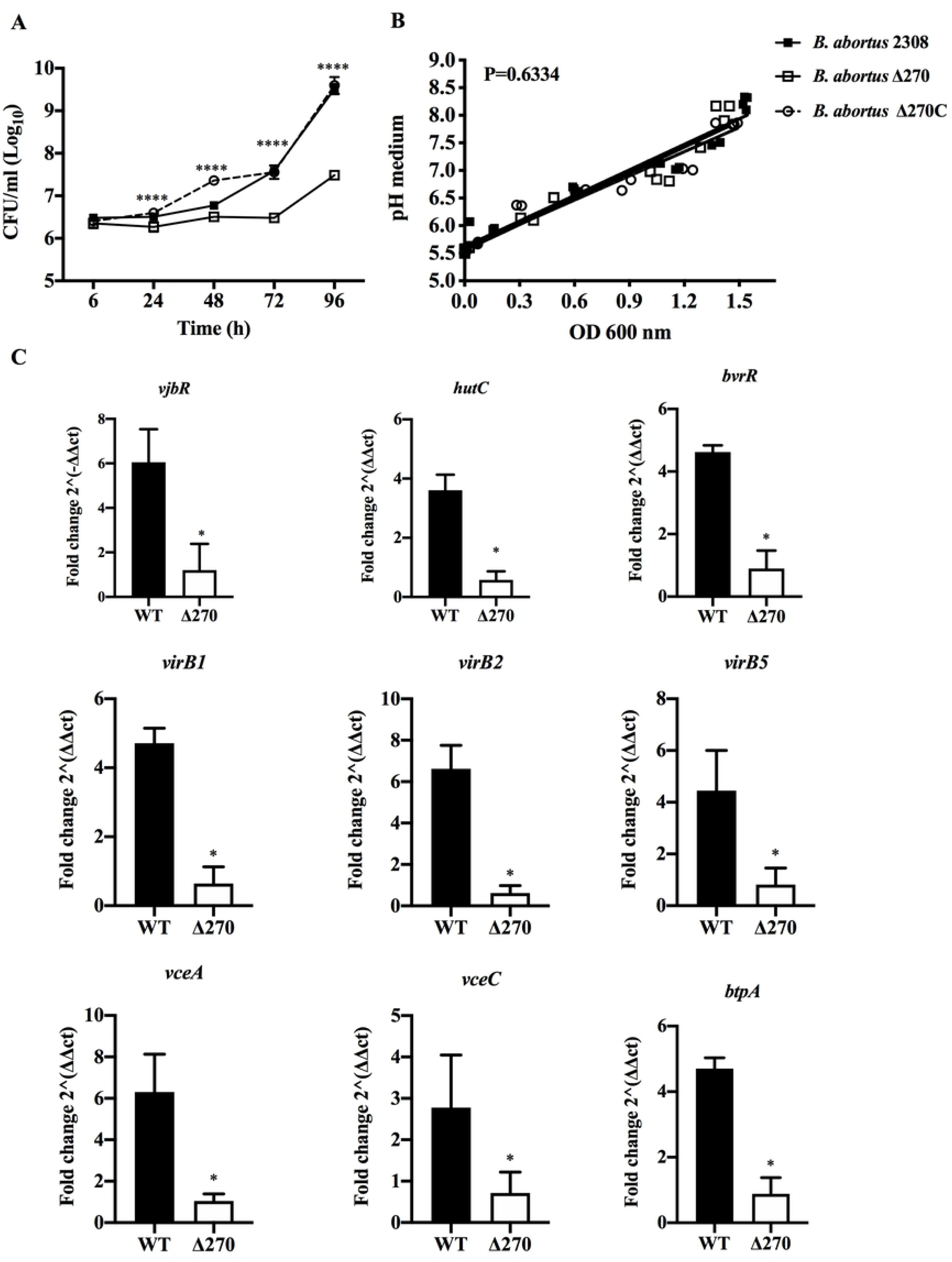
*Brucella* strains growth and gene expression under acidic stress. **A)** Growth curves of *B. abortus* 2308*, B. abortus Δ*270 y *B. abortus Δ*270C strains cultured in acidic medium (pH 5.5). **B)** Linear regression analysis of changes of pH in the medium during the growing of *Brucella* strains. **C)** The relative expression of *vjbR*, *hutC*, *bvrR*, *virB1*, *virb2*, *virB5*, *btpA*, *vceA* and *vceC* induced in *B. abortus* 2308 (WT) and *B. abortus Δ*270 cultured in medium with pH 5.5. The gene expression was calculated by 2-ΔΔ*CT* method using qPCR assays at 24 h. The housekeeping *gyrA* and *16s* genes were used as reference genes. Results were expressed as the mean ± standard deviation. *P* value < 0.05 were considered as statistically significant, where * denotes values of *P* < 0.05 and **** denotes values of *P* < 0.0001. All assays were made in triplicate.

## Discussion

*B. abortus* is a pathogen successfully adapted to survival and replicate in the intracellular environment of macrophages [4,8–11]. In these cells, it develops BCVs, which through of its interaction with the lysosomes compartments it allows to express several transcription factors regulating the expression of the T4SS virB and its effector proteins. These factors it allows to develop rBCVs, intracellularly replicate and controls the physiology of the infected cells. It was demonstrated that the ORF BAB1_0270 of *B. abortus* 2308, codifying of a Zinc-dependent metalloproteinase, also participates in the intracellular survival and capability to colonize the spleen of BALB/c mice [26]. Although the function of this Zinc-dependent metalloproteinase in the pathogenicity of this bacteria is unknown, its characterization indicating is a cytoplasmic protein that forms an operon with a transcriptional regulator containing a HTH domain of the Xre family, which could constituted a type II Toxin-Antitoxin (TA II) system, which participates in several aspects of bacterial physiology, including gene regulation, “growth arrest” and survival under environmental stress, mechanisms contributing to bacterial persistence [34,45]. Furthermore, this Zinc-dependent metalloproteinase has homology to ImmA/IrrE proteases, which cleaves repressor proteins inhibiting the expression of several genes involved in the horizontal transfer of mobile genetic elements or in the resistance to adverse conditions [31–33]. Considering this homology, we studied if this Zinc-dependent metalloproteinase of *B. abortus* would accomplish a similar function during the intracellular adaptation of *B. abortus* strain 2308 into RAW264.7 macrophages.

The infection of macrophages by *B. abortus* requires the expression of several virulence factors (e.g. Br-LPS or CβG) [12–14]. In this process, the cytoplasmic Zinc-dependent metalloproteinase participates neither the adherence nor the invasion of *B. abortus*, process where participates components from its cellular envelope such as LPS, Hsp60 or SP41 which interact with scavenger receptor SR-A, PrPc or sialic acid residues present in the macrophages lipid rafts [4,36,37]. Nevertheless, it could be involved in the ability of *B. abortus* to escape from the phagolysosomes, compartments subjecting *B. abortus* to kill by oxidative, degradative and acidic microbicides environment; and therefore, impairs it to reach the ER derived vesicles where develops rBCVs and exponentially replicate within macrophages. In order to adapt in the macrophages intracellular environment, *B. abortus* express several genes including *virB* genes (e.g. *virB1*, *virB2* or *virB5*) codifying of the T4SS, its effector proteins (e.g. VceA, VceV or BtpA) or regulators of virulence (VjbR, BvrR/BvrS or HutC) [15]. Interestingly, even when *B. abortus Δ*270 showed an up-regulation of *vjbR, hutC, bvrR, virB* during the infection of macrophages, it was insufficient to achieve a successful survival and intracellular replication. However, in the infection process the gene *vceA* codifying of a protein secreted in the cytoplasm of the macrophages and whose deletion in *B. abortus* promotes autophagy and inhibit the apoptosis in human trophoblast cells [38]. These processes are keys in the intracellular survival of this pathogen, because allows it to control the physiology of host cells. Furthermore, although the function of ORFs BAB1_ 0273 and BAB1_0627 are unknown in the physiology and/or virulence of *B. abortus*, their downregulation could indicate that the deletion of ORF BAB1_0270 affects negatively the expression of several genes potentially involved in the virulence or intracellular adaptation during the infection of macrophages.

This dysregulation in the gene expression of *B. abortus Δ*270 in an intracellular environment could have contributed to make this strain less virulent, which reduced significantly its ability to establish intracellularly and subsequently, to inhibit the expression of several proteins involved in the host’s innate and adaptive activation [39,40]. In this context, macrophages infected with *B. abortus Δ*270 were significantly stimulated, which induced the production of high levels of TNF-α, IL-6, and the expression of CD80 and CD86 co-stimulatory proteins. This process of activation could be associated to the co-localization of this mutant strain with phagolysosome cathepsin D, compartments where *B. abortus* are degraded and its antigens are recognized by pattern recognition receptors (PRR) activating of MAPK (Mitogen-Activated Protein Kinase) and/or MyD88 (Myeloid differentiation primary response 88)-dependent intracellular signaling pathways. These signaling pathways activates transcription factors such as NF-κB and Activation Protein-1 (AP-1), which promotes in macrophages the production of cytokines (IL-1β, TNF-α or IL-6), major histocompatibility complex (MHC class I and II) and the co-stimulating (CD80 and CD86) proteins activating of interferon-ã producing T CD4^+^ helper type 1 and T CD8^+^ cytotoxic, a protective response against brucellosis [23–25,40–43]. These mechanisms would demonstrate why *B. abortus Δ*270 is completely eliminated from spleen of BALB/c mice, demonstrating that the deletion of this Zinc-dependent metalloproteinase attenuate *B. abortus* 2308 and significantly reduces its biological fitness into macrophages [26].

The intracellular environment is harsh for bacteria phagocytized, because they are exposed to components such as the acidification, a factor that impair the bacterial growth and that activates bactericidal proteases in the phagosome of the macrophages [44,45]. However, it was demonstrated that *B. abortus* under pH acid express several virulence factors, including the T4SS virB to escape from phagolysosome compartments. However, our data demonstrated that deletion of this Zinc-dependent metalloproteinase significantly reduced capacity of *B. abortus* to adapt itself and grow exponentially under acidic conditions, being in addition, the gene expression of several virulence factors, including virB, effector proteins and regulators significantly down-regulated in *B. abortus Δ*270. These results may demonstrate that this Zinc-dependent metalloproteinase of *B. abortus* 2308 participates in the resistance of this bacterium to acidic stress, a condition which this bacterium is exposed during its intracellular life cycle. Therefore, is possible that, this Zinc-dependent metalloproteinase has a similar function to ImmA/IrrE proteases, which cleaves repressor proteins impeding the expression of specific genes, such as *vceA*, BAB1_0273 or BAB1_0627, which would modulate negatively the intracellular adaptation, or impeding the expression of others genes involved in the virulence of this pathogen. These observations open a spectrum of questions with respect to the function of this Zinc-dependent metalloproteinase in the virulence of *B. abortus* during the process of infection, which may be similar to that described for ImmA/IrrE proteins, that is cleaving a repressor of genes involved in the resistance to environmental stress conditions generated within phagosomes? But then, may this repressor be the transcriptional regulator present in the operon? or, is there a third element (repressor) which may be the substrate of this Zinc-dependent metalloproteinase? Besides, if we consider that this Zn-dependent metalloproteinase forms a type II TA system, then its deletion (toxin), turned the mutant strain less virulent and less resistant to intracellular microbicide mechanisms, maybe this TA system provide tolerance to stress, such as the bacterial SOS response or the production of guanosine pentaphosphate ((p)ppGpp), which contribute to bacterial persistence [34,35]. Consequently, given the importance of these TA systems, the function of this type II TA system present on the GI-3 in the physiology of *B. abortus* during its interaction with the host must be clarified. Finally, is possible that the regulatory functions associated to the type II TA system and the ImmA/IrrE-like protein participates during the intracellular adaptation of *B. abortus* in macrophages (Figure 7).

**Fig 7.**
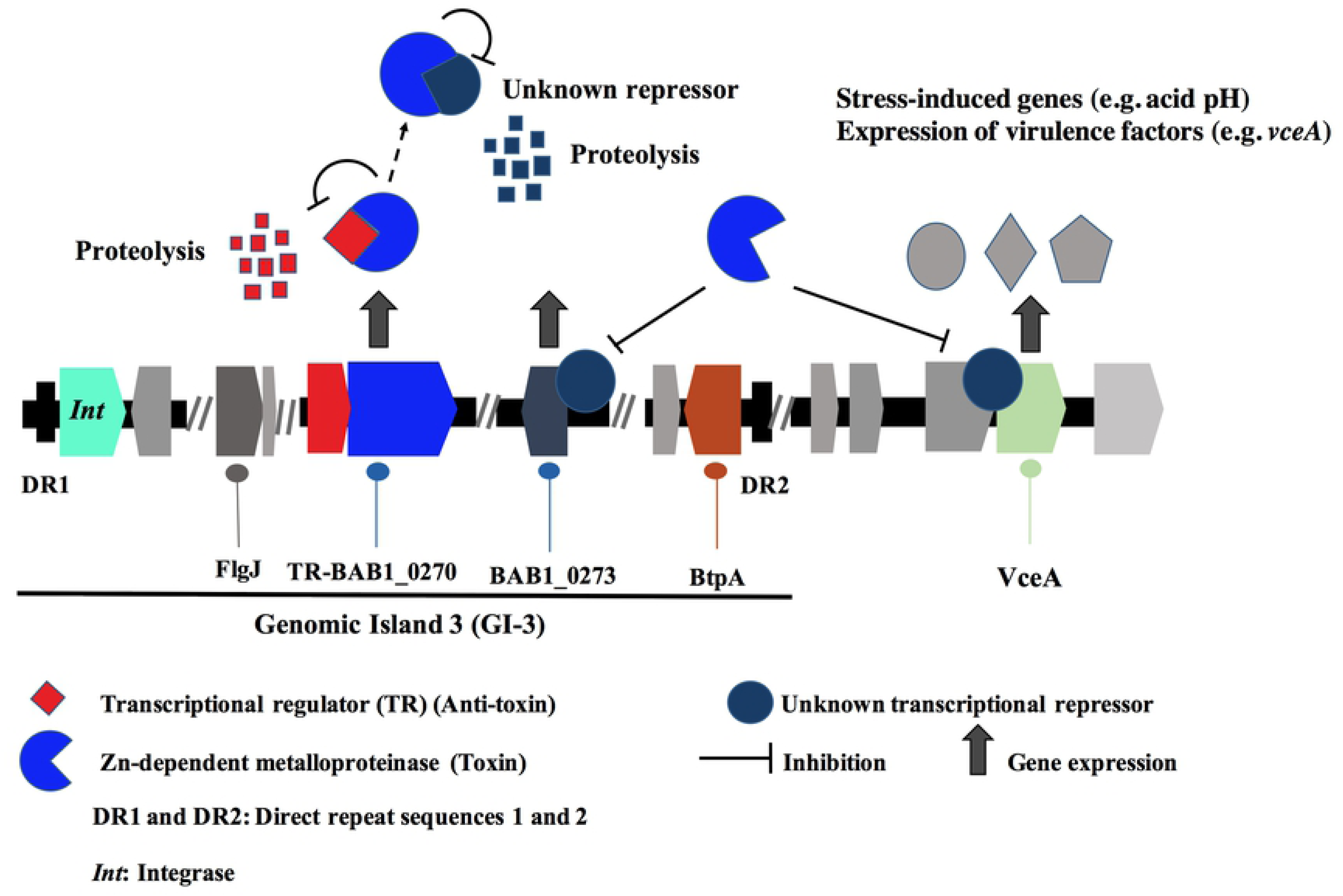
Hypothetic regulatory function of the Zn-dependent metalloproteinase. This model is based on the type II TA system and ImmA/IrrE-like regulatory functions. It postulates that operon constituted for a transcriptional regulator and Zn-dependent metalloproteinase are a type II TA system, where TR is the anti-toxin and Zn-dependent metalloproteinase is the toxin. Under stress (like acid pH) is induced the operon for bacterial persistence where anti-toxin is cleaved and Zn-dependent metalloproteinase (toxin) is free for break repressor proteins, allowing the expression of several genes involved in the bacterial resistance to hostile environment and bacterial persistence.

In summary, we postulated that this Zinc-dependent metalloproteinase not participates directly in the virulence of *B. abortus*, being its function to regulate the expression of other proteins participating in the intracellular adaptation or resistance to the microbicide mechanisms of the macrophages. Therefore, our results reported provide new insights into the mechanisms which might be regulating of the virulence of *B. abortus* during its establishment in host cells. This work provides solid evidence on how this Zinc-dependent metalloproteinase codified in the BAB1_0270 ORF contributes to the resistance of *B. abortus* 2308 during its intracellular establishment in macrophages, being its deletion the cause of a significantly reduced ability to adapt to an acidic environment, to control its endosomal traffic, to replicate intracellularly and to modulate the immune response of macrophages.

## Material and methods

### Bacterial strains and culture conditions

The strains used in this study were *Brucella abortus* 2308 (wild type, wt), *B. abortus Δ*270 (mutant for ORF BAB1_0270) and *B. abortus Δ*270C (complemented strain) [26]. For some assays, these bacteria were transformed with the broad host range vector pAKgfp1 [46] (Addgene plasmid #14076) codifying a green fluorescent protein (GFP) which allowed to study their adherence and intracellular traffic in RAW264.7 macrophages. Besides, *B. abortus* 2308 strain was inactivated using a 70% methanol-acetone solution (*B. abortus* 2308i) and used as control in some experiments. All bacteria were cultured in brucella broth (Difco) and incubated at 37°C, with agitation (120 rpm), for the time necessary according to each experiment. *B. abortus* 2308, *B. abortus Δ*270, *B. abortusΔ*270C and *B. abortus* 2308i will be hereon named WT, mutant, complemented and inactivated strains, respectively. When was necessary, broths were supplemented with antibiotics (50 μg/ml kanamycin and/or 30 μg/ml ampicillin) or their pH was adjusted to 5.5. Characteristics of bacterial strains, plasmids and primers used in this work are described in the Table 1. Primers for mutant strain are described in the **S1 Table**. All the assays and experiments were done following the procedures established by the Biosafety Committee of the University of Concepción.

**Table 1.** Bacterial strains, plasmids and primers used for mutant strains.

| Bacterial strains | Characteristics | Reference |
| --- | --- | --- |
| *B. abortus*2308 | Wild type, smooth and virulent strain | Laboratory stock |
| *B. abortus* Δ270 | *B. abortus* 2308 mutant for BAB1_0270 ORF (Km<sup>r</sup>) | [26] |
| *B. abortus* Δ270C | *B. abortus* Δ270 (Km<sup>r</sup>) complemented with pBV1 codifying of BAB1_0270 ORF | [26] |
| *B. abortus* 2308-*gfp* | Wild type, smooth and virulent strain transformed with pAKgfp1 plasmid codifying of green fluorescence protein (GFP), (Amp<sup>r</sup>). | This work |
| *B. abortus* Δ270-*gfp* | *B. abortus* 2308 mutant for BAB1_0270 ORF (Km<sup>r</sup>) and transformed with pAKgfp1 plasmid codifying of green fluorescence protein (GFP) (Amp<sup>r</sup>). | This work |
| *B. abortus* 2308i | Wild type strain inactivated with 70% of methanol-acetone. | This work |
| *Escherichia coli* BL21 (DE3) | Strain used for recombinant protein production. DE3 is a recombinant phage harboring the T7 RNA polymerase gene. | [55] |

| Plasmids | Characteristics |  |
| --- | --- | --- |
| pSIM7/pSIM9 | Broad-host-range cloning vector (Cm<sup>r</sup>), Lambda Red Recombinase (λ-Red). | [56] |
| pKD4 | Cassette for Km<sup>r</sup> sequence | [57] |
| pVB1 | Cloning vector for PCR products (Amp<sup>r</sup>) | [26] |
| pVB1-BAB1_0270 | Recombinant plasmid (Amp<sup>r</sup>) containing the BAB1_0270 ORF. | [26] |
| pAKgfp1 | Broad-host-range cloning vector pBBR1MCS4 (Amp<sup>r</sup>) codifying of green fluorescence protein (GFP). | [46] |
| pColdII | Vector derived from backbone plasmid pUC118 is used for cold shock-induced protein expression in *E. coli* followed by a 6xHis tag. | [58] |

### Cell line and culture conditions

RAW264.7 murine macrophages (American Type Culture Collection, ATCC) were seeded in 12 or 24 wells Nunclon Delta Surface plates (ThermoFisher Scientific, MA, USA) containing Dulbecco’s Modified Eagle Medium (DMEM) supplemented with 10% fetal bovine serum (FCS) plus 100 UI penicillin, 100 μg/ml streptomycin and 0.25 μg/ml amphotericin B (Antibiotic-antimycotic solution, ThermoFisher Scientific). Cells were incubated at 37°C in a 5% CO_2_ environment for the necessary time in accordance with the experiments described below.

### *In silico* analysis of Zinc-dependent metalloproteinase structure

The physicochemical parameters of this Zn-dependent metalloproteinase, such as amino acid composition, molecular weight (MW), theoretical isoelectric point (pI), extinction coefficient, instability index, *in vitro* and *in vivo* half-life, grand average of hydropathicity (GRAVY) and aliphatic index were evaluated with ProtParam server at http://web.expasy.org/protparam [47]. Their antigenicity was analyzed using the VaxiJen v2.0 server (http://www.ddgpharmfac.net/vaxijen/VaxiJen/VaxiJen.html), which employs an alignment-independent method based on auto cross covariance (ACC) [48]. Then, a homology modeling of the hypothetical protein was done using I-TASSER (Iterative threading assembly refinement) server http://zhanglab.ccmb.med.umich.edu/I-TASSER/ [49], which employs a hierarchical approach to predict the protein structure based on multiple-threading alignments and fragment assembly simulations methods. I-Tasser expresses the confidence of the modeling as a C-score that varies from −5 to 2, where the high C-score is related to high confidence. The visualization of the 3D structures was done with PyMOL. In order to refine the whole hypothetical protein, the best model structure obtained from I-TASSER was introduced to the GalaxyRefine server at http://galaxy.seoklab.org/cgi-bin/submit.cgi?type=REFINE. This server uses mild and aggressive relaxation methods by reconstruction of side chains [50]. Later, the final refinement of the model was used for energy minimization with KobaMIN server at http://csb.stanford.edu/kobamin/ [51]. In order to recognize the potential errors in the initial and final tridimensional structure of the hypothetical protein, the PDB formats were charged in ProSA-web server at https://prosa.services.came.sbg.ac.at/prosa.php [52]. ProSA-web is based on energy distribution and comparison with the native similar structure of a database. The quality of the 3D predicted protein is shown as a Z-score, that indicates possible errors when it is compared with native experimentally determined protein stored in a database.

### Purification and characterization of the Zn-dependent metalloproteinase

This metalloproteinase is codified in the ORF BAB1_0270 of *B. abortus*, which was expressed cloning its DNA sequence (GenBankAM040264.1) in the vector of expression pColdII (pColdII-Zn-dMP). This recombinant vector was used to transform *Escherichia coli* BL21(DE3) strain, where the recombinant protein was purified. Visualization of this recombinant protein was analyzed by Western blot, using antibodies anti-6xHis Tag (Abcam, Cambridge, UK). Subcellular location, promoter and conserved domains, were analyzed using bioinformatic predictions by CELLO v2.5 [53] (http://cello.life.nctu.edu.tw/), BPROM-prediction for bacterial promoters [54], and protein-protein Basic Local Alignment Search Tool (BLASTp), respectively. Because these metalloproteinases contain COG2856 domains, which usually are associated to putative operons with proteins containing Helix-turn-Helix (HTH) domains of the Xre family, a search was conducted to identify this transcriptional regulator in the *B. abortus* 2308 and determine if both proteins are part of an operon. These analyses were confirmed extracting genomic DNA (gDNA) using a Wizard® Genomic DNA Purification Kit (Promega, WI, USA) and total RNA was extracted with TRIzol (ThermoFisher Scientific Inc, MA, USA) as indicated by the manufacturer. cDNA was obtained from total RNA by RT-PCR using Maxima First Strand cDNA Synthesis Kit (ThermoFisher Scientific Inc, MA, USA) and the expression of the BAB1_0270-Transcriptional regulator operon was analyzed by PCR-amplification of both genes from genomic DNA (gDNA) and cDNA, using specific primers (Table 2). PCR products were visualized by agarose gel electrophoresis at 1%.

**Table 2.**
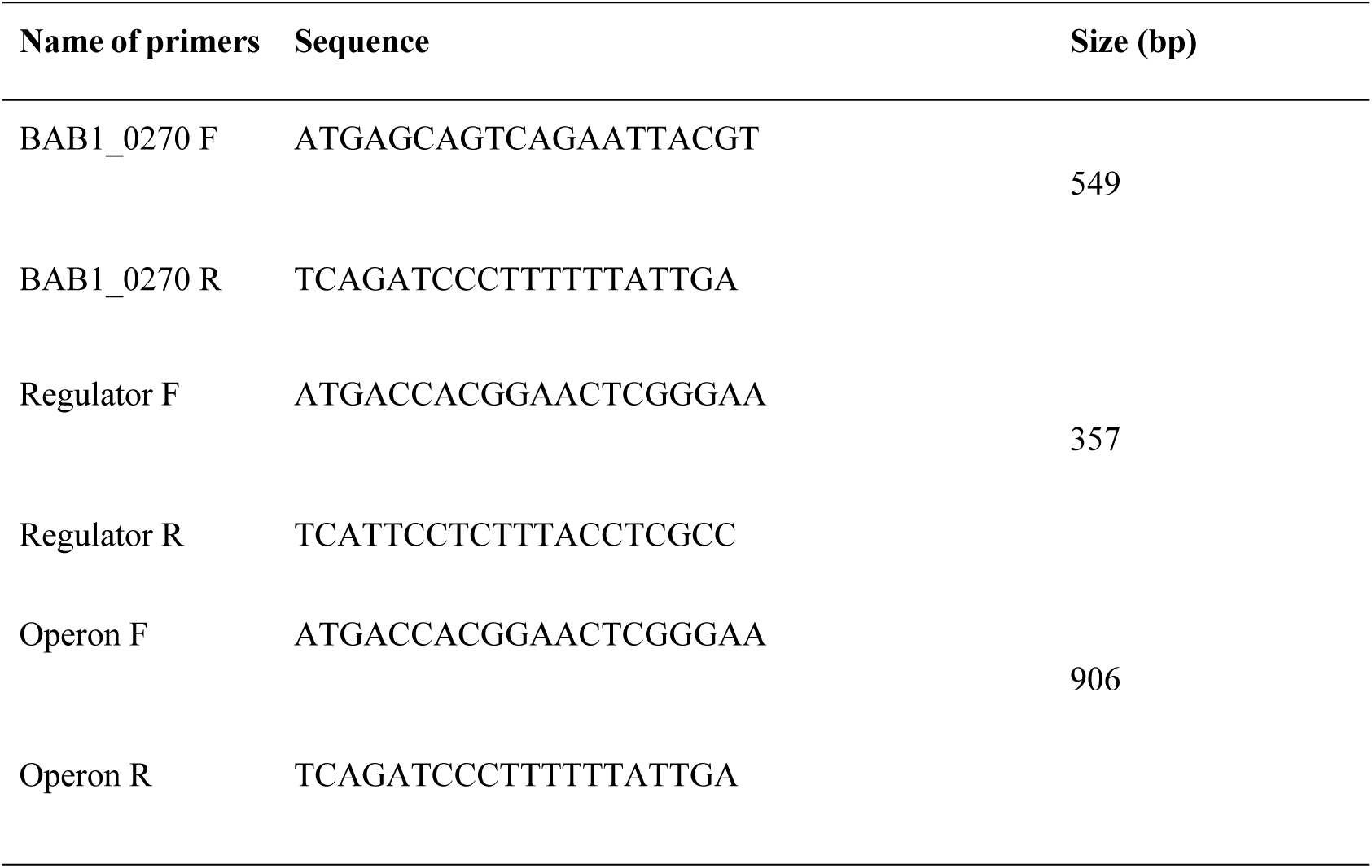
Primers used in this study for operon identification

### Adhesion assays

The role of ORF BAB1_0270 in the adherence of *B. abortus* strains to RAW264.7 macrophages was examined using strains *B. abortus* 2308-GFP, *B. abortus Δ*270-GFP and *B. abortus Δ*270C-GFP. Bacterial adherence was analyzed by culturing 1 × 10^5^ RAW 264.7 macrophages adhered to glass coverslips (Thermo Fisher Scientific Inc., MA, USA) and treated with 0.5 mg/ml cytochalasin D, an inhibitor of phagocytosis, to impair bacterial internalization. Then, cells were infected with *Brucella*-GFP strains at a multiplicity of infection (MOI) of 1:10 for 30 min at 37°C in 5% CO_2_. Next, the cells were washed using PBS, fixed with 4% paraformaldehyde (PFA) and permeabilized with cold methanol (−20°C) for 10 seconds. Actin filaments were labeled with phalloidin Alexa Fluor 633 (Thermo Fisher Scientific Inc., MA, USA) diluted at 1:500 for 1 h at 37°C. Finally, the samples were mounted on slides using Dako Cytomation fluorescent mounting medium (Dako North America, Inc., USA). All samples were observed using a Zeiss LSM 700 laser scanning confocal microscope (Zeiss, Oberkochen, Germany). Images were acquired and assembled with the IMARIS software from data obtained from three independent experiments.

### Intracellular trafficking

The contribution of the BAB1_0270 ORF to the intracellular traffic was evaluated by co-localization of *B. abortus* 2308-GFP, *B. abortus Δ*270-GFP and *B. abortus Δ*270C-GFP with Early Endosome Antigen 1 (EEA1, an early endosomal protein), cathepsin D (a phagolysosomal protein) and calnexin (an endoplasmic reticulum protein) using confocal microscopy. For this, macrophages where adhered to coverslips and inoculated with different *B. abortus*-GFP strains at a MOI 1:10 during 5 min, 10 min and 15 min for EEA1, and 1 h and 12 h for cathepsin D and calnexin. Post-infection (pi), macrophages were fixed with 4% PFA, permeabilized with 0.1% Triton X-100 and incubated with goat anti-EEA1 polyclonal (Santa Cruz Biotechnology, Dallas, TX, USA), goat anti-cathepsin D (Abcam, Cambridge, UK) or rabbit anti-calnexin (Abcam, Cambridge, UK) antibodies for 3 h in a humidity chamber. All antibodies were diluted in PBS (7.4 pH) supplemented with 0.5% bovine serum albumin (BSA). After incubation, the coverslips were washed using PBS (pH 7.4) and incubated with donkey anti-goat IgG Alexa Fluor 594 (Thermo Fisher Scientific Inc., MA, USA) or donkey anti-rabbit IgG Alexa Fluor 647 as secondary antibody (Abcam, Cambridge, UK) diluted 1:500. Finally, the samples were mounted on slides using Dako Cytomation fluorescent mounting medium (Sigma-Aldrich, St. Louis, MO, USA). All samples were observed using a Zeiss LSM 700 laser scanning confocal microscope (Zeiss, Oberkochen, Germany). Images were acquired and assembled with the ImageJ software from data obtained (percentages of co-localization) of two independent experiments.

## Intracellular survival of *B. abortus* into macrophages

The role of the BAB1_0270 ORF in the intracellular survival of *B. abortus* 2308 was studied infecting RAW264.7 macrophages with wt, mutant and complemented strains. For this, 2.5 × 10^5^ RAW264.7 macrophages were seeded per well in 24-well plates (Nunclon Delta, Denmark) containing incomplete DMEM medium and incubated for 2 h at 37°C to allow for adherence. In parallel, *B. abortus* strains, cultured for 48 h at 37°C, were harvested in the logarithmic phase by centrifugation (2000 xg for 10 min), resuspended in incomplete DMEM medium at 2.5 × 10^6^ colony forming units (CFU)/ml and added to the adhered macrophages at MOI 1:10. Then, plates were centrifuged at 500 x g for 10 min to facilitate contact between bacteria and macrophages and incubated for 1 h at 37°C in 5% CO_2_. Next, extracellular bacteria were eliminated by replacing the incomplete medium by DMEM medium supplemented with 10% FCS plus 100 μg/ml gentamycin and 50 μg/ml streptomycin. Infected cells were incubated during 6 h and 24 h at 37°C in 5% CO_2_. The intracellular survival of the *B. abortus* strains was evaluated collecting the cells in phosphate buffer saline-EDTA (PBS, pH 7.4; EDTA 2 mM) by centrifugation (160 x g for 10 min) and lysing them with PBS-Tritón X-100 (0.1%). Bacterial counts were determined by plating serial dilutions of bacteria in Brucella agar for 72 h at 37°C. All assays were done in triplicate.

## Production of TNF-α and IL-6

The contribution of the ORF BAB1_0270 in the inflammatory response of macrophages infected with *B. abortus* was assessed through the production of the pro-inflammatory cytokines TNF-α and IL-6 by ELISA. For this, 5 × 10^5^ RAW264.7 macrophages/well were seeded in 24-well plates (Nunclon Delta, Denmark) and infected with wt, mutant and inactivated strains at MOI 1:10 under the culture conditions described above. The production of these cytokines was quantified in the supernatants from macrophages infected 6 h and 24 h pi. TNF-α and IL-6 production was evaluated by ELISA, using the commercial kits eBioscience Mouse TNF alpha and Mouse IL-6 ELISA Ready-SET-Go! (Fisher Scientific, MA, USA). Final concentrations of cytokines were quantified by standard curves based on the concentration of recombinant mouse TNF-α and IL-6. Results were obtained using a VictorX3 ELISA reader (PerkinElmer, Waltham, MA, USA) at 450 nm. All assays were done in triplicate.

## Flow cytometry

The contribution of the ORF BAB1_0270 on the expression of co-stimulatory proteins was evaluated in infected macrophages by flow cytometry. For this, 5 × 10^5^ RAW264.7 macrophages/well were seeded in 24-well plates (Nunclon Delta, Denmark) and infected with wt, mutant and inactivated strains at MOI 1:10 under the culture conditions described above. Control groups with non-infected (non-stimulated) and lipopolysaccharide (LPS) stimulated (5ìg/ml) (LPS from *E. coli* O26:B6 strain, Sigma-aldrich) macrophages were included in the experimental design. At 6 and 24 h pi, macrophages were collected from the wells, washed using PBS-EDTA and incubated with Zombie Violet (BioLegend, CA, USA) for viability during 30 min. Cells were washed using PBS-EDTA-BSA and stained with anti-CD11b (clone M1/70) antibodies conjugated to APC/Cy7, anti-CD80 (clone 16-10A1) conjugated to FITC and anti-CD86 (clone GL-1) conjugated to APC (BioLegend, CA, USA) diluted 1:500. Samples were incubated for 30 min in darkness, washed and fixed with 0.5% paraformaldehyde. For data acquisition, cells were resuspended in PBS and analyzed in a BD LSR Fortessa X-20 (BD Biosciences) flow cytometer. The acquired data was analyzed using FlowJo (BD Biosciences). All assays were done in triplicate.

## RT-qPCR assays

The relative expression of several genes codifying of T4SS virB, its effectors protein and transcription factors involved in the virulence and intracellular survival of *B. abortus Δ*270 was evaluated in infected macrophages by 2^−ΔΔ*CT*^ method. For this, 1 × 10^6^ RAW264.7 macrophages/well were infected at MOI 1:10 for 24 h with the strains wt, mutant and inactivated. Then, total RNA was extracted with TRIzol (ThermoFisher Scientific Inc, MA, USA) as was indicated by the manufacturer. Complementary DNA (cDNA) was obtained from RNA by reverse transcription using the Maxima First Strand cDNA Synthesis kit for RT-PCR (ThermoFisher Scientific Inc, MA, USA) and the relative expression of the genes of interest (Table 3) was quantified using the Takyon q-PCR kit for SYBR assays by means of the AriaMx Real Time PCR system (Agilent Technologies, CA, USA). *gyrA* and *16s* housekeeping genes were used as reference genes for all assays. All assays were done in triplicate.

**Table 3.**
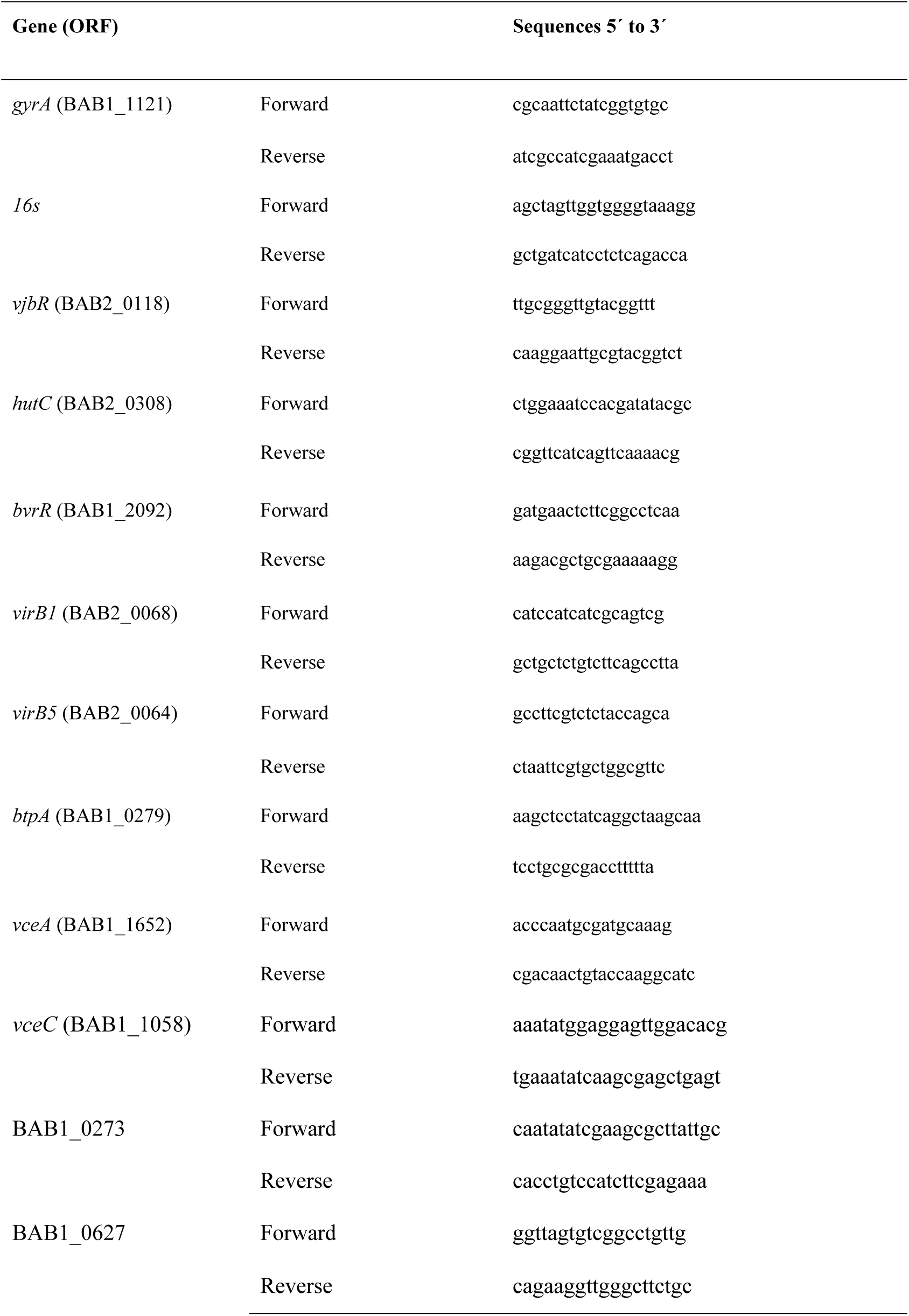
Primers used in this study for qPCR assays.

## Growth and gene expression of *B. abortus* in acidic stress

The contribution of the ORF BAB1_0270 to resistance of *B. abortus* 2308 to acidic stress was evaluated culturing wt, mutant and complemented strains in 50 ml of Brucella broth adjusted to pH 5.5. Bacteria were cultured at 37°C with agitation (120 rpm) during 96 h. At times 6, 24, 48, 72 and 96 h, 1 ml aliquots were obtained to count bacteria. Serial dilutions were plated on Brucella agar for 72 h and results were reported as CFU/ml. In parallel, pH of supernatant from *B. abortus* strains culture was measured using a digital pH-meter. In addition, relative expression of genes involved in the virulence of *B. abortus* cultured for 24 h in medium with pH 5.5 (see primers in Table 3) was evaluated by 2^−ΔΔ*CT*^ method using the RT-qPCR assays described above. For these assays, mRNA from *B. abortus* 2308 cultured under physiological pH was used for calibration of gene expression, and *gyrA* and *16s* housekeeping genes were used as reference genes for all assays. All assays were done in triplicate.

## Statistical analysis

Data obtained from the adherence and intracellular co-localization experiments were analyzed by means of a T-test. Intracellular survival of bacteria, cytokines, co-stimulatory proteins and bacterial growth under acidic stress conditions were analyzed by means of a two-way ANOVA. Changes in the pH by *B. abortus* strains were measured using a simply linear regression. The gene expression was analyzed by Non parametric Mann-Whitney U Test. All analyses were done using the GraphPad Prism 8 software. Results were expressed as the mean ± standard deviation. Values of *P* < 0.05 were considered as statistically significant.

## Acknowledgements

This work was supported by Grant 1180122 from the Fondo Nacional de Desarrollo Científico y Tecnológico (FONDECYT), Santiago, Chile. Leonardo A. Gómez is a recipient of a CONICYT scholarship for PhD students in Chile.

## Author Contributions

**Conceptualization:** Leonardo A. Gómez, Angel A. Oñate.

**Data Curation:** Leonardo A. Gómez, Francisco I. Álvarez, Angel A. Oñate.

**Formal analysis:** Angel A. Oñate.

**Founding acquisition:** Angel A. Oñate.

**Methodology:** Leonardo A. Gómez, Francisco I. Álvarez, Raúl Molina, Rodrigo Soto, Carla Daza-Castro, Manuel Flores, Yrvin León, Angel A. Oñate.

**Supervision:** Angel A. Oñate.

**Writing – Original draft:** Leonardo A. Gómez, Angel A. Oñate.

**Writing – review & editing:** Leonardo A. Gómez, Francisco I. Álvarez, Yrvin León, Angel A. Oñate.

